# Decision signals in the absence of spiking activity in primate visual cortex

**DOI:** 10.1101/2025.07.29.667496

**Authors:** Yueyue S. Hou, Liu D. Liu, Christopher C. Pack

## Abstract

Fluctuations in single-neuron activity in sensory cortex often correlate with perceptual decisions. This kind of correlation has been hypothesized to reflect a causal influence of sensory signals on decisions, a feedback influence of decisions on sensory signals, and various other factors as well. To disentangle these different possibilities, we have examined local field potentials (LFPs) recorded from the middle temporal (MT) area of non-human primates performing a motion discrimination task. Compared to single-neuron spiking, LFPs have the advantage of being decomposable into different frequencies that are associated with different anatomical sources of input. More importantly, they persist when spiking activity is inactivated, which precludes a causal influence of the corresponding neural activity on behavior. We found that high frequency (70–150 Hz) LFP power was correlated with perceptual decisions and that this correlation disappeared when spikes were inactivated, consistent with a causal role for this frequency band in decision making. These signals overlapped in time with decision signals in lower gamma-band power (30–70 Hz), which persisted after spiking inactivation, suggesting a non-causal, feedback input. Interestingly, lower-frequency LFP signals (5–30 Hz) reflected both impending perceptual decisions and the outcome of previous trials. Our results therefore reveal that neural activity multiplexes different sources of information about perceptual decisions and that these types of information can be estimated reliably from different LFP frequencies.

## Introduction

Many previous studies have attempted to quantify the relationship between spiking activity and perceptual decisions ^1–4^. In these studies, the degree to which single-neuron activity predicts a subject’s decision is often quantified with a metric called choice probability (CP). Despite its widespread use, the interpretation of CP is still debated ^5^. CP was originally assumed to represent a feedforward, or causal, influence of neural activity on decisions ^1^, while subsequent work favored a feedback influence of decision-making processes on neural activity ^6^. A third alternative is that another factor, such as reward ^7^ or motor planning ^8^, influences both neural activity and decision-making.

Under reasonable assumptions ^9–11^, one can infer the importance of these different influences by examining the dynamics of CP over time ^6,12,13^. For instance, feedback is often assumed to be slower than feedforward processing, so that it is more prominent during the later phases of the neuronal response to a stimulus. However, this is not always the case ^14^, and spike timing can be affected by many factors that are not necessarily related to decisions ^14,15^. Given the correlational nature of CP, disentangling the various contributions to decision-related signals remains a significant challenge in the field.

As an alternative to single neurons, one can examine decision-related signals in local field potentials (LFPs), which reflect the coordinated activity of neural populations ^16,17^. LFPs are characterized in terms of oscillations, and there is evidence that higher-frequency LFP oscillations are predominantly linked to feedforward sensory encoding ^18–22^, while lower-frequency LFPs primarily represent feedback processing ^18,21–23^. Thus, examining CP in LFPs across specific frequency bands could provide information about the composite nature of CP signals.

To investigate this possibility, we recorded spikes and LFPs from the middle temporal (MT) area in two macaque monkeys performing a random dot motion discrimination task. This allowed us to compute CP in different LFP frequency bands and to provisionally estimate the properties of feedforward and feedback contributions to CP. To test these estimates more directly, we temporarily silenced local spiking activity pharmacologically, thereby eliminating the possibility of a feedforward influence of MT activity on perceptual decision-making.

We found that significant CP signals were present in the high-gamma frequency range (70– 150 Hz), suggesting a feedforward influence. Consistent with this idea, inactivation of spiking activity abolished CP in this frequency band. In contrast, CP in the low gamma frequency band (30–70 Hz) persisted despite inactivation of spiking activity, suggesting that these CP signals were not causal in nature. CP detected in alpha and beta frequency bands (5–30 Hz) appeared to reflect a third type of influence, which was largely attributable to the outcome of previous trials. Our findings thus demonstrate that CP reflects multiple decision-related signals, which can be effectively distinguished through LFPs in different frequency bands.

## Results

We trained two macaque monkeys to discriminate the coherent motion of a random dot kinematogram (RDK), while we recorded and manipulated neural activity in visual cortical area MT (Fig. 1A) ^4^. As illustrated in Figure 1B, the task required monkeys to fixate a central point on the screen while a motion stimulus was displayed within the receptive fields of concurrently recorded MT neurons. Following a fixed delay, the monkeys reported the stimulus motion direction by making a saccade toward one of two targets aligned with the axis of motion, located on opposite sides of the initial fixation point. Task difficulty was titrated by varying dot coherence randomly from trial to trial ^4^.

**Figure 1.**
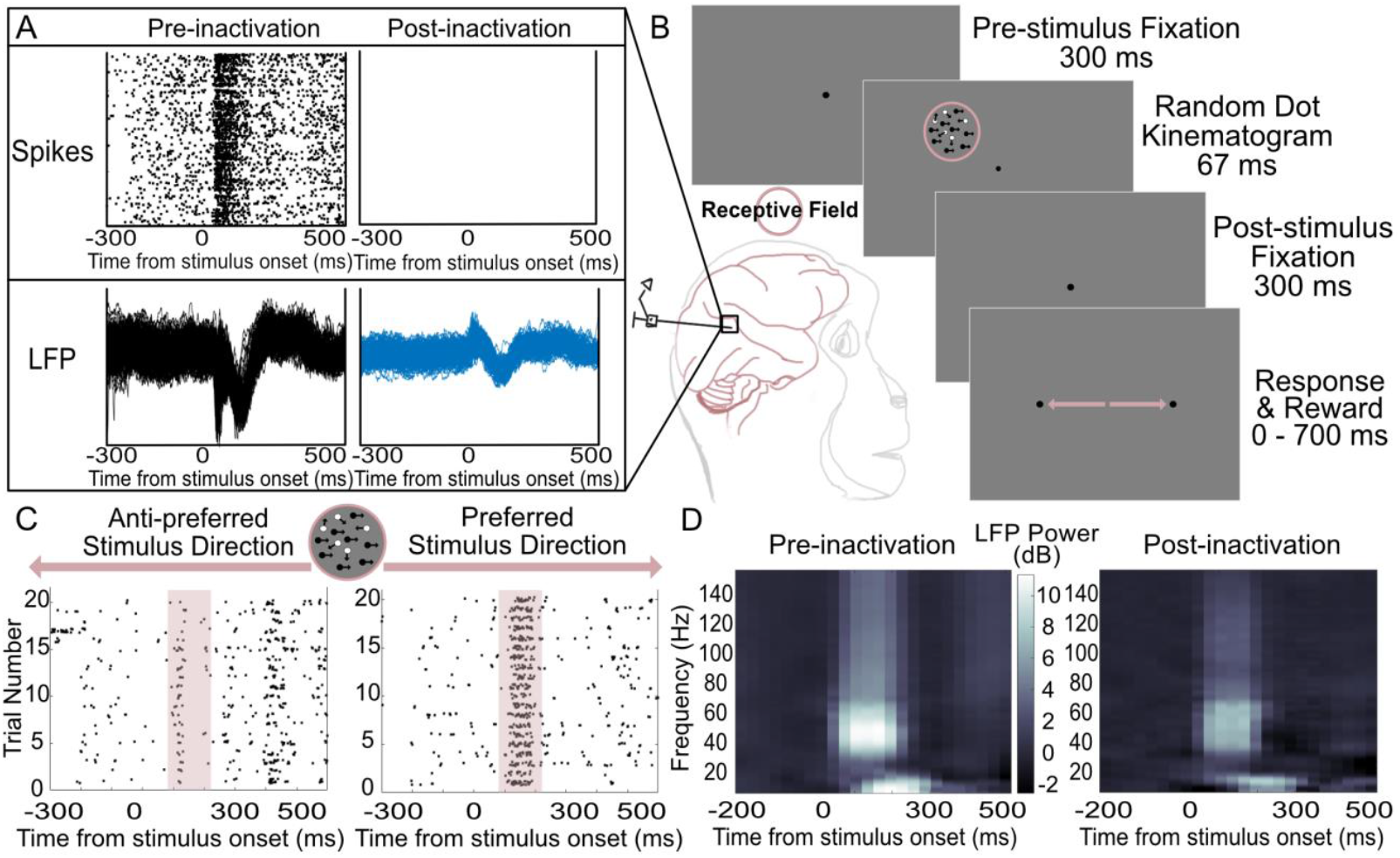
Measuring and calculating local field potential (LFP)-based choice probability (CP). **A)** A 16-channel linear electrode array was used to simultaneously record multiple neurons’ spiking activity and LFPs. Within each recording session, the GABA_a_ agonist muscimol was used to temporarily inactivate the middle temporal (MT) region to eliminate local spiking activity; LFP signals could still be detected. **B)** During each recording session, two rhesus monkeys performed a random dot motion discrimination task. The animals were trained to distinguish whether dots were moving in the neurons’ preferred or anti-preferred stimulus direction. The stimulus was placed in the receptive field (pink circle) of the MT neurons. Task difficulty was adjusted by varying the percentage of dots moving coherently in one direction (filled circles), while other dots moved at random (open circles). Note that the dot types, pink-colored circle and arrows are for illustration only and were not shown to the subjects. **C)** Recorded neurons’ direction selectivity was determined by multi-unit spike counts within the stimulus response epoch before muscimol injection (shaded panels). **D)** Time-frequency analysis of LFP before and after muscimol inactivation in area MT. The color shows LFP power normalized to the baseline, as a function of time relative to stimulus onset and of frequencies between 0 and 150 Hz exclusively.

On each trial, the stimulus motion was in the neurons’ preferred or anti-preferred direction. We defined the preferred direction at each recording site based on multi-unit spike counts (Fig. 1C), which exhibit greater direction selectivity than LFP power ^17,24^ (see *Methods* for details).

### Decision Information in the LFP

CP quantifies the likelihood that an ideal observer could predict each trial’s perceptual decision from concomitant fluctuations in neural activity ^4^. Although it is typically computed from spike rates, CP can also be computed from trial-to-trial variations in LFP power ^16,17^. In this case, values above 0.5 indicate higher LFP power on trials in which behavioral choices aligned with the recorded neurons’ preferred stimulus direction; values below 0.5 indicate an inverse relationship between LFP power and preferred-direction choices. Due to the non-normal distribution of LFP power, we computed CP in the LFPs using robust scaling—normalization based on median and interquartile range—rather than the pooled z-scoring method that is commonly used for spike data ^25^. We validated this approach through simulations (see *Methods* and *Supplementary Materials*).

In each session, we computed LFP-based CP before and during inactivation of spiking activity with the GABA agonist muscimol. Although muscimol consistently reduced spiking rates to zero ^26^, LFP activity persisted (Fig. 1D), suggesting that it contains information about input to the MT recording sites ^24^. Indeed, following muscimol inactivation, LFP power in response to the visual stimuli decreased on average by 76.0%, with no consistent difference across frequencies (Spearman’s ρ = 0.067, permutation test, P = 0.418). Because the CP metric is sensitive to behavioral performance ^27,28^, we equalized overall performance by selecting different RDK coherence levels for the pre- and post-muscimol trials (see *Methods*).

### CP as a Function of LFP Frequency and Behavioral Epoch

The spectrograms in Figure 2A show average CP values as a function of LFP frequency and time relative to stimulus onset, based on eight sessions in two animals. As indicated by the brownish colors in the figure, CP values above 0.5 arose for a restricted range of times and LFP frequencies. Figure 2B shows that the administration of muscimol altered the distribution of CP across time and frequency.

**Figure 2.**
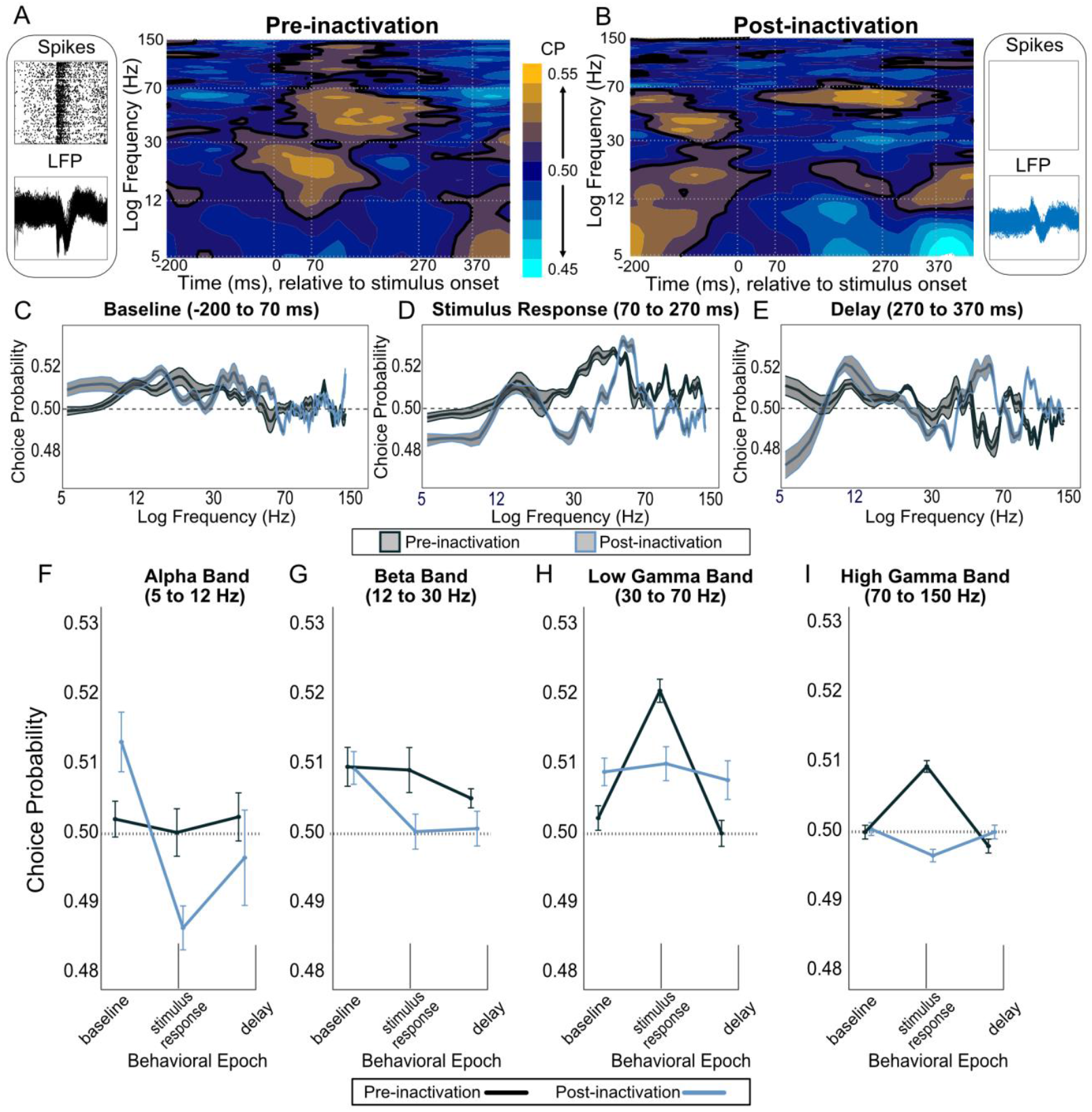
LFP-based decision signals before and after muscimol inactivation as a function of frequency and time. **A)** CP heatmap generated from LFP power when local spiking activity was active. The color gradient bar indicates CP values at chance level (navy blue), above chance level (yellow), and below chancel level (cyan). Vertical gray dotted lines (from left to right) indicate the stimulus onset, stimulus offset, and the duration of post-stimulus fixation period that covers both stimulus response and delay epochs. The outlined contour (black) indicates CP values equal to and above 0.51. A first-order Savitzky-Golay filter was applied to smooth the signal and reduce distortion of overall trends. **B)** CP heatmap in the absence of spiking activity after muscimol inactivation. **C-E)** CPs as a function of LFP frequency across three behavioral epochs (**C)** baseline, **D)** stimulus response, and **E)** delay). Standard errors were denoted by colored ribbons with black as no inactivation and cerulean as inactivation. **F-I)** LFP-based CPs across task epoch (baseline, stimulus response, and delay) and frequency bands: **F)** alpha (5*–*12 Hz), **G)** beta (12*–*30 Hz), **H)** low gamma (30– 70 Hz), and **I)** high gamma (70*–*150 Hz). Black curves correspond to the aggregated values before muscimol inactivation. Cerulean curves indicate aggregated data after muscimol inactivation. The horizontal dotted line at the chance level was drawn in each subpanel. The error bars denote ± standard error of the mean (SEM).

To estimate the most relevant LFP frequencies, Figures 2C–E present the same data averaged within task-relevant time periods: the baseline (−200 to 70 ms), covering the pre-stimulus fixation; the stimulus response (70 to 270 ms); and the delay (270 to 370 ms) before the appearance of the choice targets. CP peaked for different frequencies across the three time periods. These dynamics were well captured by collapsing the LFP frequencies into canonical bands (Fig. 2F–I): alpha (5 to 12 Hz ^29^), beta (12 to 30 Hz ^17,30^), low gamma (30 to 70 Hz ^31–33^), and high gamma (70 to 150 Hz ^30,34^). We next examined these frequency bands in detail.

### High Gamma Choice Probability is Consistent with a Feedforward Influence

For high gamma oscillations, CP peaked in the stimulus response epoch (Figure 2I), with the average CP value being significantly above chance (0.510 ± 0.008, P < 0.001, Wilcoxon signed-rank (WSR) test). Following muscimol injection, the mean CP value at an equivalent behavioral performance level declined significantly (0.497 ± 0.008, P < 0.001, WSR test; P < 0.001, Wilcoxon rank-sum (WRS) test); these findings were consistent across individual monkeys (Supplementary Fig. 1; Panel A, Monkey Y: no inactivation, CP = 0.510 ± 0.012, P < 0.001, WSR test; with inactivation, CP = 0.497 ± 0.009, P < 0.001, WSR test; P < 0.001, WRS test; Panel B, Monkey C: no inactivation, CP = 0.507 ± 0.013, P < 0.001, WSR test; with inactivation, CP = 0.497 ± 0.011, P = 0.002,WSR test; P < 0.001, WRS test). The reduction in CP after inactivation was largely restricted to the stimulus response period (Fig. 2A–B, D, I), which is consistent with previous studies indicating that high gamma oscillations often reflect the propagation of sensory signals to higher cortical areas as a feedforward influence ^21,22,29,35^.

### Low Gamma Choice Probability Reflects Correlated Decision Information

Low-gamma (30–70 Hz) choice signals also appeared in MT during the stimulus response epoch (Fig. 2H), with an average CP value of 0.521 ± 0.010, significantly higher than 0.5 (P < 0.001, WSR test). Within this time period, low-gamma CP peaked at 120 ms, compared to 80 ms for high-gamma CP. The delayed emergence of CP in the lower gamma band is consistent with the idea that some component of CP reflects feedback from higher-level cortical areas ^6^.

Indeed, choice signals in the low-gamma band persisted after muscimol inactivation, with the average CP value being 0.510 ± 0.015, which was significantly above chance (P < 0.001, WSR test). The results were consistent across two monkeys (Supplementary Fig. 1; Panel A, Monkey Y: no inactivation, CP = 0.531 ± 0.017, P < 0.001, WSR test; with inactivation, CP = 0.523 ± 0.019, P < 0.001, WSR test; P < 0.001, WRS test; Panel B, Monkey C: no inactivation, CP = 0.509 ± 0.016, P < 0.001, WSR test; with inactivation, CP = 0.502 ± 0.017, P = 0.443, WSR test; P < 0.001, WRS test). Thus, the continued presence of low gamma decision signals—despite the elimination of local spiking— strongly suggests a component of CP that is not causal in nature.

Further insight into the nature of this component is given by the timing of the post-inactivation CP signal. Following inactivation, low gamma CP became most prominent at the end of the stimulus response period, around 260 ms (Fig. 2B), peaking in Monkey Y at 240 ms and in Monkey C at 280 ms. This was not due to a delay in stimulus processing, as time-frequency power analyses revealed only a general reduction in the evoked LFP magnitude after muscimol (Fig. 1D). Instead, this temporal shift likely reflects reliance on memory-driven or other feedback mechanisms during the post-stimulus fixation period ^11,36–38^.

### Alpha-Beta Involvement and Reward History

Decision-related signals in the alpha-beta band (5–30 Hz) were different between the two monkeys. In Monkey Y, average CP values during the stimulus period were significantly above chance in the beta range (12–30 Hz), both before and after muscimol inactivation (Supplementary Fig.1A; no inactivation: CP = 0.522 ± 0.021, P < 0.001; with inactivation: CP = 0.503 ± 0.024, P < 0.001; WSR test). In contrast, Monkey C showed below-chance CP values in the same frequency range across both conditions (Supplementary Fig.1B; no inactivation: CP = 0.491 ± 0.015, P < 0.001; with inactivation: CP = 0.488 ± 0.019, P < 0.001; WSR test). The persistence of CP sign following inactivation in both monkeys is consistent with previous findings indicating that beta LFPs are less associated with feedforward encoding ^21,29^.

Indeed, we observed above-chance CPs in the alpha and beta frequency ranges even during the pre-stimulus baseline period (see Fig. 2C, F–G). Before inactivation, this choice-related activity differed between animals; Monkey Y exhibited significant beta-band CPs (Supplementary Fig. 1A: CP = 0.523 ± 0.019, P < 0.001, WSR test), while Monkey C did not (Supplementary Fig. 1B: CP = 0.497 ± 0.013, P = 0.025, WSR test). Nevertheless, pre-stimulus CPs after inactivation were consistent across monkeys (Supplementary Fig. 1; Panel A, Monkey Y: CP = 0.508 ± 0.017, P < 0.001; Panel B, Monkey C: CP = 0.516 ± 0.015, P < 0.001; WSR test). This individual-level discrepancy prior to the stimulus presentation suggests differences in internal states or task strategies between monkeys ^39,40^.

One possibility suggested by previous studies is that CP on one trial could be influenced by the reward (or lack thereof) on the previous trial; this could affect arousal, which in turn could modulate both neural responses and behavioral performance ^7,41,42^. In that case, CP during the baseline period in the alpha and beta bands might capture this covariation between neural and behavioral signals, without necessarily being related to the decision *per se*. To assess this possibility, we separated CP for trials that were preceded by a successful (rewarded) trial and those that had been preceded by an unsuccessful (non-rewarded) trial (see *Methods*).

In both monkeys, we found that CP in the alpha and beta frequencies was higher after a non-rewarded trial than after a rewarded trial (Fig. 3A, C). Before inactivation, alpha-beta CPs following a failed trial averaged 0.517 ± 0.027 (P < 0.001, WSR test) for Monkey Y, peaking in the beta band (Fig. 3A), and 0.513 ± 0.023 (P < 0.001, WSR test) for Monkey C, peaking in the alpha band (Fig. 3C). In contrast, alpha-beta CPs following a successful trial were significantly lower, averaging 0.508 ± 0.019 (P < 0.001, WSR test) for Monkey Y (Fig. 3A) and 0.496 ± 0.007 (P < 0.001, WSR test) for Monkey C (Fig. 3C), with no significant peak found in any frequency band.

**Figure 3.**
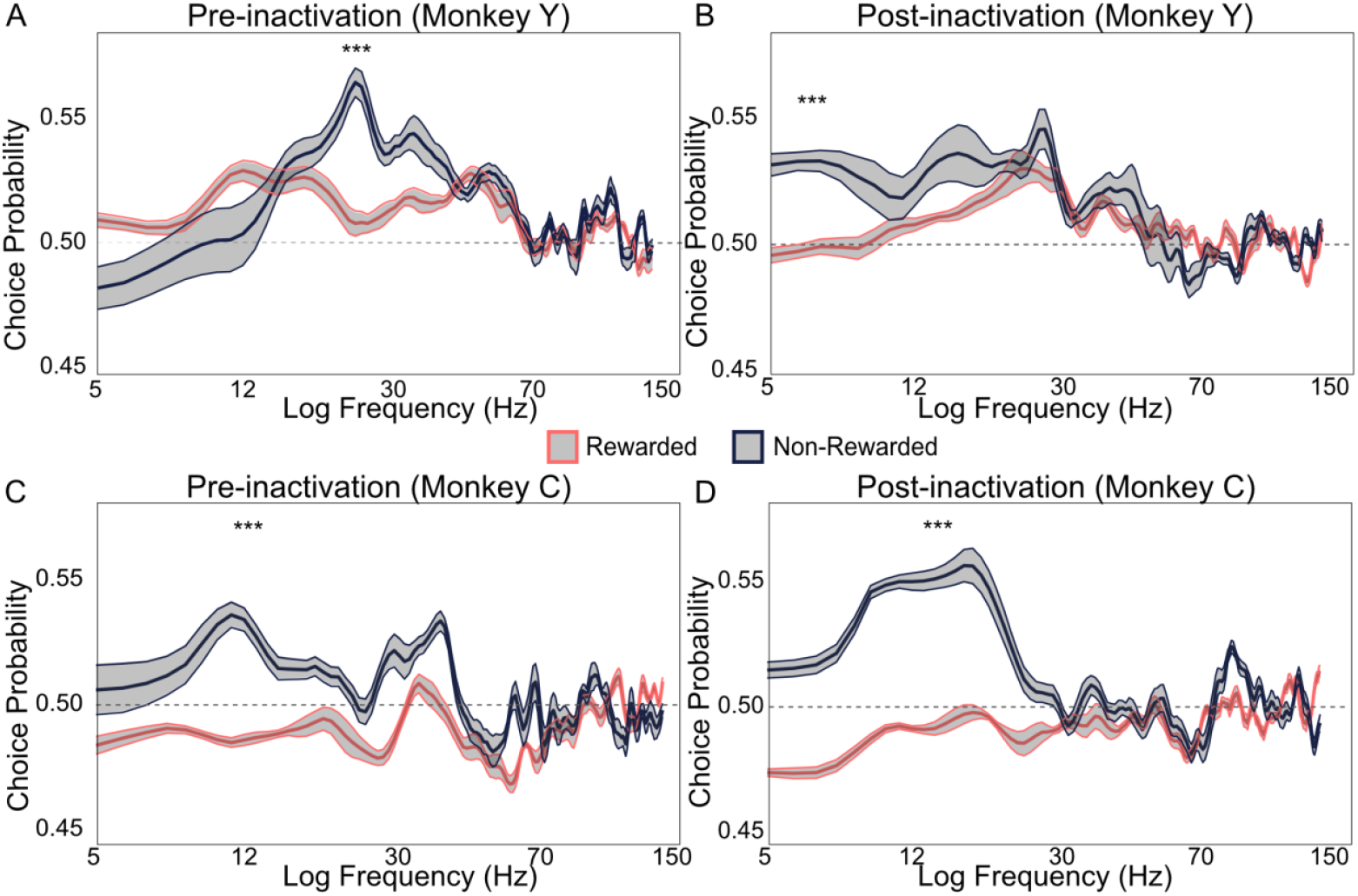
Alpha-beta CPs during the baseline period for individual monkeys. **A) and B)** Monkey Y; **C) and D)** Monkey C. **A) and C)** show CPs as a function of frequency before inactivation, while **B) and D)** show CPs after inactivation. Red curves represent conditions in which previous trials were rewarded; black curves represent conditions in which previous trials were non-rewarded. Horizontal dashed gray lines indicate chance-level CP. Shaded ribbons show SEM; asterisks denote statistically significant differences between rewarded and non-rewarded conditions.

This pre-stimulus component of CP following an unsuccessful trial was not reduced after muscimol inactivation (WRS test; Monkey Y, P = 0.367; Monkey C, P = 0.391; Fig. 3B, D). Interestingly, the between-monkey difference in baseline was reflected in successful trials, where above-chance CPs for rewarded trials after inactivation were found only in Monkey Y (Fig. 3B; CP = 0.523 ± 0.028, P < 0.001, WSR test) and not in Monkey C (Fig. 3D; CP = 0.489 ± 0.012, P < 0.001, WSR test), implying that Monkey Y was overall more influenced by preceding reward outcomes. These findings suggest that pre-stimulus CP may reflect non-sensory signals that nevertheless have an influence on behavior ^7,40,43^.

## Discussion

Using LFP-based CP measurements and causal inactivation in non-human primates performing a visual task (Fig. 1), we have identified distinct frequency-dependent features of decision-related neural activity. High gamma frequencies appear to reflect feedforward decision signals, as they disappeared with muscimol inactivation (Fig. 2). In contrast, decision signals in the low gamma frequency persisted with inactivation, indicating that this component of CP did not reflect a direct causal influence (Fig. 2). CP in alpha and beta bands revealed influences that likely reflect modulation based on reward history (Fig. 3).

### Comparison with previous work

Extensive research has used CP to investigate the neural bases of decision-making across various brain regions, including the parietal cortex ^13,28,44^ and early visual areas ^39,45,46^, using behavioral paradigms such as visual discrimination ^3,4,47^ and change-detection tasks ^48,49^. Recent work in this area has hypothesized different time courses for feedforward and feedback influences on CP ^6,11^. Feedforward components of CP are thought to arise early in the stimulus response period, with feedback influences arising later ^6,9,50,51^. These correlational studies are, however, limited by the fact that the relationship between the timing of signals and their anatomical source is not entirely straightforward in visual cortex ^14^. Moreover, fluctuations in neural activity are not strictly attributable to task performance ^8,26,44^. Thus, there is increasing emphasis on causal methods for probing the circuitry of decision-making ^45,51^.

For example, inactivating parts of parietal cortex, which exhibits strong choice-related signals, does not necessarily affect behavioral performance, revealing a non-causal role for these signals ^13,44,52,53^. In contrast, choice-related activity in sensory areas like MT might reflect a more causal influence, as inactivation of these areas significantly impairs performance on perceptual tasks ^13,44,52^. Given the known functional coupling between MT and LIP, it is conceivable that decision-related signals in LIP are fed back to MT, leading to correlated activity even in the absence of a causal sensory role ^45^. Indeed, feedback projections are known to influence sensory processing and attention in MT, though their functional role in decision-making is unclear ^54,55^. We recently showed that CP in MT can also reflect oculomotor signals ^8^.

A few previous studies have attempted to evaluate CP based on LFP power ^16,17^. In one study, the authors recorded spikes and LFPs from V1 and V4 while monkeys performed a two-alternative forced-choice orientation discrimination task ^16^. Similar to our findings, they reported decision-related information in the alpha and beta frequencies, as well as in high-gamma frequencies during the stimulus response period. Liu and Newsome recorded LFPs from area MT during a speed discrimination task and found significant LFP-based CP in frequencies above 40 Hz, while LFP power below 30 Hz actually exhibited below-chance CP values ^17^. It is difficult to compare these studies directly, given substantial differences in behavioral tasks and analytical methods, but the general conclusion that CP can be computed from LFP appears to be robust.

### Frequency-specific neural computations in decision-making

The strong reduction of high gamma CP signals following muscimol inactivation (Fig. 2) supports the hypothesis that these signals represent feedforward processing ^20,21,56^. This might result from the fact that higher-frequency LFPs are generally found in the upper layers of cortex, which send feedforward projections to higher-level areas ^30^, although we were unable to verify the laminar locations of our recordings.

The persistence of low-gamma CP signals after muscimol inactivation could reflect feedback signals from decision-related areas ^6,9,50^. Alternatively, they could reflect shared noise correlations between neural activity in MT and other areas, such as V3A ^57^ or MST ^58^. These could be feedforward in nature, in the sense that their sensory activity might inform decisions, rather than the other way around. Evidence in favor of a feedback source for low-gamma CPs comes from the fact that they occurred later in time than the high-gamma CPs. Moreover, after inactivation, the peak low-gamma CP shifted into the delay period, consistent with previous findings that working memory influences low gamma oscillations in MT ^36,37^.

CP signals in the alpha and beta bands were also found before and after inactivation (Fig. 3). Given that the strongest effects were observed during the pre-stimulus baseline, this component of CP cannot be attributed to feedback about the decision. Rather, they seem to reflect the reward outcome of previous trials, as has been found for spiking activity ^7^.

One interpretation of this finding is that reward transiently boosts arousal or attention, which influences both stimulus coding and task performance ^7,40^.

## Conclusions

Much previous work has been concerned with the question of whether decision-related signals in visual cortex are feedforward or feedback in nature. By analyzing LFP signals in different frequency bands, we detected both sources of decision-related activity in visual cortex, with largely overlapping time-courses. We also detected a third source related to the outcome of previous trials. The relative balance of these signals likely depends on the structure of the behavioral task and the strategies used by the subjects. Given this complexity, our results suggest that LFPs can be useful in revealing information about perceptual decision-making that is not readily apparent in spiking activity.

## Materials and Methods

The experimental methods were detailed in previous papers ^8,26,59^; this paper primarily involves a re-analysis of the associated data.

### Observers and Ethics Statement

Two adult female rhesus macaque monkeys (*Macaca mulatta*) (Monkey Y, age: 10 years; Monkey C, age: 8 years; both weighted 7 kg) participated in this study. The animals were housed in a temperature- and pressure-controlled socially accessible environment under a 12-hour light/dark cycle. Behavioral training and experimental recordings were conducted during the light phase of the cycle. Animals sat comfortably while head-fixed in a custom-designed primate chair. Identical stimuli, timing, and rewards were used for both monkeys. A solenoid-operated reward system was used to dispense juice reward to the monkeys (Crist Instruments Co., Inc).

All procedures adhered to the regulations established by the Canadian Council on Animal Care and were approved by the Institutional Animal Care Committee of the Montreal Neurological Institute’s Animal Care Committee (protocol no. 5031).

### Experimental Setup

Prior to the experiments, an MRI-compatible titanium head post (Hybex Innovations, Montreal) and a plastic recording cylinder were affixed to each monkey’s skull under general anesthesia. Area MT was identified based on transitions from white matter to gray matter, as well as the electrode depth, the prevalence of robust, direction-selective visual responses, and the relationship between receptive field size and eccentricity. Eye movements were monitored with an infrared eye tracking system (EyeLink1000, SR Research) with a sampling rate of 1,000 Hz.

### Neural Recording

Single units were recorded utilizing linear microelectrode arrays (V-Probe, Plexon) comprising 16 contacts. Initially, the electrode array was lowered to the designated depth, followed by the estimation of multi-channel receptive fields through the manual positioning of a moving bar within the visual field. Visual motion stimuli were presented at a frequency of 60 Hz with a resolution of 1,280 by 800 pixels. The viewing area covered 60° by 40° at a distance of 50 cm. Neuronal signals were continuously monitored during acquisition via computer display. Spike signals were thresholded in real-time, with local field potential signals undergoing band-pass filtering at 0.5 Hz to 150 Hz, while spike signals were band-pass filtered at 150 Hz to 8 kHz, with monitoring conducted on an oscilloscope and loudspeaker. Spike signals were thresholded in real time, and spikes were assigned to single units by a template-matching algorithm (Plexon MAP System). Then, spikes were manually sorted offline using a combination of automated template matching, visual inspection of waveforms, clustering in the space defined by the principal components, and absolute refractory period (1 ms) violations (Plexon Offline Sorter).

### Visual Stimuli, Training, and Behavioral Paradigm

The monkeys were trained to perform a motion discrimination task, with random dot kinematograms (RDK). During each session, neuronal direction and speed preferences were measured using 100% coherent dot patches positioned within the receptive fields found at the recording site on that day. The receptive field locations were quantified by fitting a 2D spatial Gaussian to the neuronal response measured across a 5 × 5 grid of stimulus positions in an offline analysis. The grid comprised of moving dot patches centered on the initially hand-mapped receptive field locations. We confirmed that all neurons included in our analysis had receptive field centers within the stimulus patch used for the behavioral experiments. The speed of the dots was chosen to match the preference of the MT neurons in proximity to the recording site. The direction of motion was consistently selected based on the preferred or anti-preferred direction of the neurons being examined. The stimulus size also matched the receptive field size of the MT neurons (mean radius = 6.3° ± 1.2°). The RDK coherence was randomly selected on each trial from seven values that encompassed the range of the monkey’s psychophysical threshold (2%, 4%, 8%, 16%, 32%, 64%, 100%).

The individual trial structure of a random dot motion discrimination task is detailed below (also see Fig. 1B). On each trial, animals established and maintained fixation for a 300 ms, followed by a brief presentation of the dot coherence patch (typically for 67 ms) on the receptive field centers. The monkeys were then required to maintain fixation for another 300 ms, after which the fixation point disappeared, two choice targets appeared, and the monkey made a saccade to the corresponding target to indicate its perceived motion direction (preferred or anti-preferred relative to the isolated neurons) in exchange for juice rewards. Targets were typically positioned at 10° eccentricity, with each pair of angles sampled at 45° intervals on the screen. For example, if a neuron’s preferred direction was 45°, the targets were intentionally positioned to correspond with both the preferred and anti-preferred directions, meaning they would be placed at 45° and 225°, respectively. The distance between the two saccade targets was typically 20°. The appropriate saccade direction correlated with the most similar saccade target direction (i.e., the monkey performed the rightward saccade to correctly report rightward motion). The monkey was required to indicate its decision within 700 ms after the onset of the choice targets. In trials with no motion signal (0% coherence), rewards were randomly assigned in half of the instances. If fixation was disrupted at any time during the stimulus, the trial was aborted. In a typical experimental session, the monkeys performed the task on 20 to 40 repetitions of each stimulus.

### Infusion of Muscimol

The linear V-Probe contained a glass capillary with an inner diameter of 40 μm. The capillary was positioned at one end between contacts 5 and 6 of the array, with contact 16 located at the most dorsal-posterior position. The opposite end was connected via polyethylene tubing (PE20, inner diameter = 0.38 mm) to a 10 μl micro-syringe (Hamilton) to ensure continuous and smooth injection.

Muscimol, a GABA_a_ agonist (Sigma), was dissolved in sterilized saline (pH ∼= 7) to achieve a concentration of about 10 mg/ml ^60^. In half of the sessions, muscimol (generally 2 μl at a rate of 0.05 μl/min) was administered via the fluid channel in the V-Probe to inhibit its adjacent neural activity.

Behavioral testing commenced 40-50 minutes post-infusion at each site. The probes remained at the infusion depth within the cortex for the duration of the session, allowing us to verify that spiking activity remained silenced throughout the post-muscimol period, which typically lasted 2.5 to 3.5 hours. The inactivated area returned to its normal state typically within two days. Experiments involving muscimol were performed at a frequency not exceeding once per week.

As describe previously ^26^, there was no behavioral effect of injecting saline or of injecting muscimol at sites distant from the recording site, indicating that the muscimol results were due to local inactivation, rather than to any general effects of the injection.

### Analyses of Neural and Behavioral Responses

Only recording sessions that contained pre- and post-muscimol data within the same recording session were included for analysis. We excluded trials in which monkeys failed to execute a saccade toward one of the choice targets or broke fixation during the stimulus presentation or delay periods. Overall, a total of 5 control sessions and 5 corresponding inactivation sessions from monkey Y were analyzed, along with 3 control sessions and 3 corresponding inactivation sessions from monkey C. On average, monkey Y completed 318 trials per session, and monkey C performed 358 trials per session, encompassing both correct and error trials of the task. The same number of trials were performed after the infusion of muscimol.

#### LFP preprocessing and re-referencing

Local Field Potentials (LFPs) from the 16th channel, located at the most dorsal-posterior position, were utilized as the reference signal, while the LFP signals from the other channels were re-referenced accordingly so that artifacts from eye blinks and other physiological factors were eliminated. The line noise at 60 Hz was removed in LFP raw signals. We utilized the Chronux toolbox ^61^ in MATLAB, applying its multi-taper method for Fourier power spectral estimation, using two tapers derived from discrete prolate Slepian sequences to achieve a frequency resolution of 1 Hz.

To compare LFP power before and after muscimol injection, we used a non-parametric statistical method. First, we calculated the log ratio of LFP power spectra (post-vs. pre-inactivation) across trials from all sessions. The difference in total power before and after inactivation was quantified by computing the area under the Power Spectral Density (PSD) curves (MATLAB’s *trapz* function). To assess whether this power difference varied significantly as a function of frequency, we computed Spearman’s rank correlation coefficient (ρ), testing statistical significance through a permutation test (n = 5,000 permutations).

#### LFP-based choice probabilities (CP)

The computation of LFP-based CP, a metric that quantifies trial-to-trial fluctuations between LFP power variability and psychophysical choices, was conducted, mostly using established methods developed for spiking activity^4^.

We first subdivided trials into two choices defined by neurons’ direction selectivity and behavioral outcomes. We included sites for which the preferred direction had at least a 50% higher multi-unit spike count in the stimulus response period (70 – 270 ms relative to stimulus onset) than the spikes in the baseline (−200 – 0 ms relative to stimulus onset). The stimulus response interval was selected due to the significant correlation between the spikes observed during this time window and the animals’ behavioral choices ^59^. Direction selectivity was assessed across multiple neurons at the highest dot coherence levels (64% and 100%) for every electrode channel independently.

In this way, a preferred choice contained trials that exhibited either a preferred direction with a correct response or a non-preferred direction with an incorrect response; an anti-preferred choice contained trials that involved either a non-preferred direction with a correct response or a preferred direction with an incorrect response.

Because CP is meant to be insensitive to the mean amplitude of neural responses, most analyses of CP first involve a normalization of firing rate, using z-scoring or balanced z-scoring ^25^. However, given the non-normal distribution of LFP power, this approach has the potential to introduce biased estimate of CP. To assess this possibility, we used the NeuroDSP toolbox ^62^ to simulate LFP data from 100 trials of one-second duration (Supplementary Fig. 2A), each comprised of stochastic periodic and aperiodic components; these were generated using the function *sim_combined* with the default parameters. We implemented this procedure twice to simulate trials associated with two behavioral choices that differed in mean amplitude across frequencies, one being 100 times greater in amplitude than the other (Supplementary Fig. 2A). Then, we computed spectrograms to represent the simulated LFP data before normalization (Supplementary Fig. 2B). A proper normalization would yield CP values of 0.5, because CP should be insensitive to the mean of the neural signals. However, we found that z-scoring and balanced z-scoring failed to eliminate the mean difference between the signals associated with the two choices, largely distorting the signals that were associated with lower LFP power (Supplementary Fig. 2C). This distortion arises because in non-Gaussian distributions like LFP power, the smaller absolute values in the low-power group make it more susceptible to noise. Thus, z-scoring disproportionately emphasizes outliers when the distribution is non-normal, biasing LFP-based CP estimation.

We therefore used a robust scaling method that does not rely on any specific distributional assumptions but rather quantifies the central tendency and dispersion by median and interquartile range. In our simulated example, robust scaling aligned the two choices to a uniform scale with identical values, thereby recovering the accurate CP values across all frequencies (Supplementary Fig. 2D). This simple simulation confirmed that robust scaling effectively returned CPs to chance, demonstrating its suitability for normalizing LFP power.

Following appropriate normalization, the distributions of two normalized choices were subsequently analyzed using receiver operating characteristic (ROC) analysis to compute CP. CP was determined by calculating the area under the ROC curve. The area under the ROC curve ranges from 0.0 to 1.0, reflecting the performance of an ideal observer in determining motion direction based on the trial-to-trial LFP power. Values of 1.0 and 0.0 represent perfect classification, whereas a value of 0.5 signifies performance equivalent to random chance classification.

CP can be sensitive to overall behavioral performance, and the infusion of muscimol significantly impaired the monkeys’ performance. It was therefore necessary to equalize performance pre- and post-muscimol by selecting trials with different stimulus coherence levels. For Monkey Y, we used coherence levels of 4% and 8% before muscimol injection and 8%, 12%, and 16% after injection. For Monkey C, we used coherence levels of 4%, 8%, and 12% before muscimol injection and 8%, 12%, and 16% after injection. These stimulus levels equated behavioral performance before and after inactivation, at approximately 65% correctness across monkeys (Monkey Y, 66%; Monkey C, 62%).

We then estimated CPs based on LFP power. First, we calculated the CP for each frequency bin individually, utilizing power measurements obtained from a sliding window of 200 ms width, advancing in 20 ms increments. We calculated the averaged CPs across three distinct epochs relative to stimulus onset: baseline (−200 to 70 ms), which occurred between the pre-stimulus fixation and stimulus presentation, stimulus response (70 to 270 ms), which spanned between stimulus offset and post-stimulus fixation, and delay (270 to 370 ms) spanning before the appearance of the choice targets. We statistically quantified by averaging power across four frequency bands: alpha (5 to 12 Hz), beta (12 to 30 Hz), low gamma (30 to 70 Hz), and high gamma (70 to 150 Hz).

To examine how reward history affects pre-stimulus CP (Fig. 3), we subcategorized CP estimates into rewarded and non-rewarded conditions based on their preceding trials’ behavioral outcome. The rest of the computation was identical to that descried above.

### Quantification and Statistical Analyses

We used nonparametric techniques for the statistical evaluation of our data. All statistical studies were performed using MATLAB’s core tools and custom programs. We employed the Wilcoxon signed-rank (WSR) test to assess the significance level of LFP-derived CPs relative to 0.5. The difference in LFP-derived CPs before and after muscimol infusion was evaluated using the Wilcoxon rank-sum (WRS) test. Significance was determined at p < 0.05.

## Supplementary Materials

**Supplementary Figure 1.**
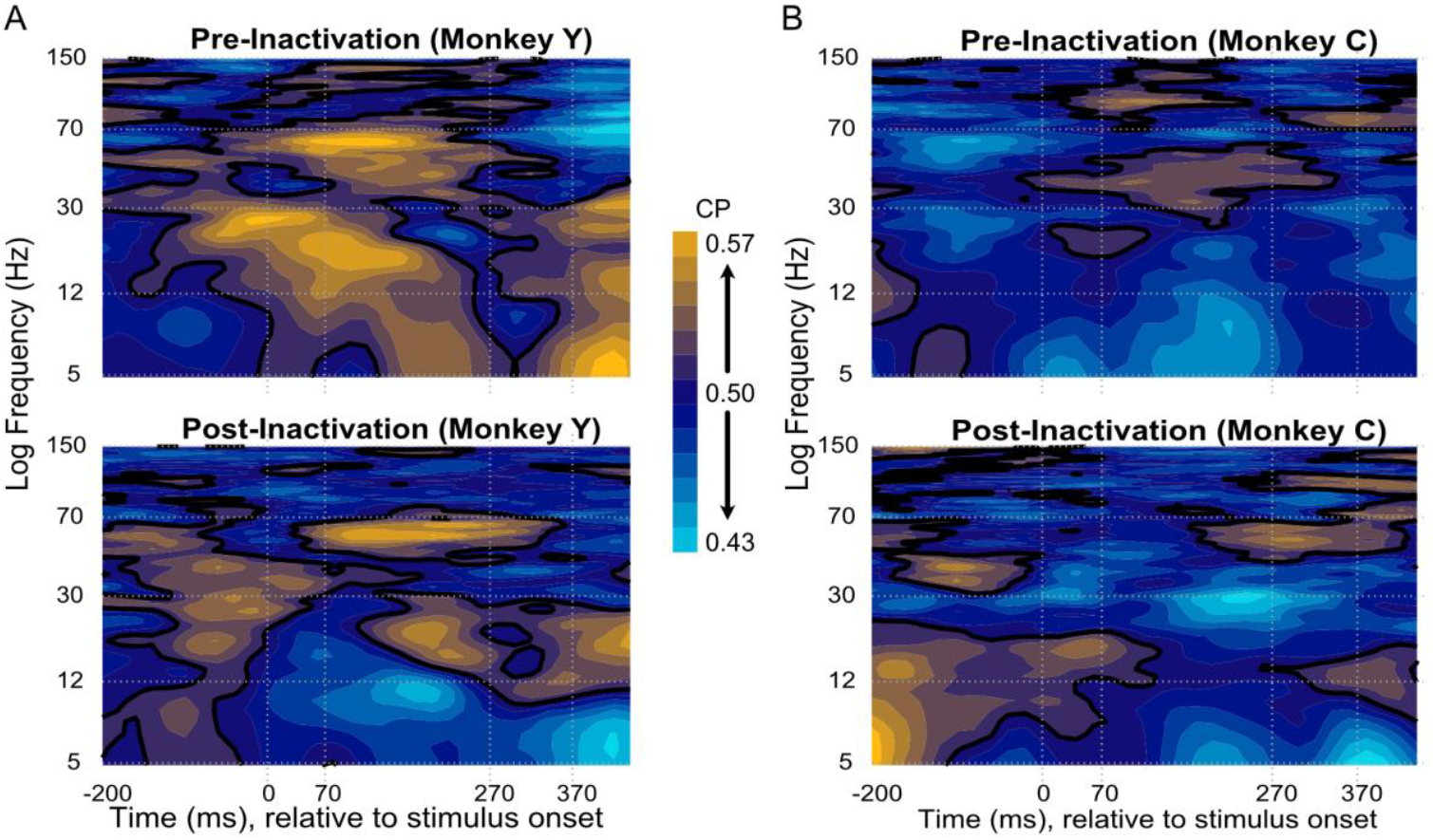
LFP-based CPs of **A)** monkey Y and **B)** monkey C before muscimol inactivation (upper panel) and after muscimol inactivation (lower panel). The panels show LFP-based CP levels as a function of log frequency (Hz) and time aligned to stimulus onset (ms) as a continuous heatmap.

**Supplementary Figure 2.**
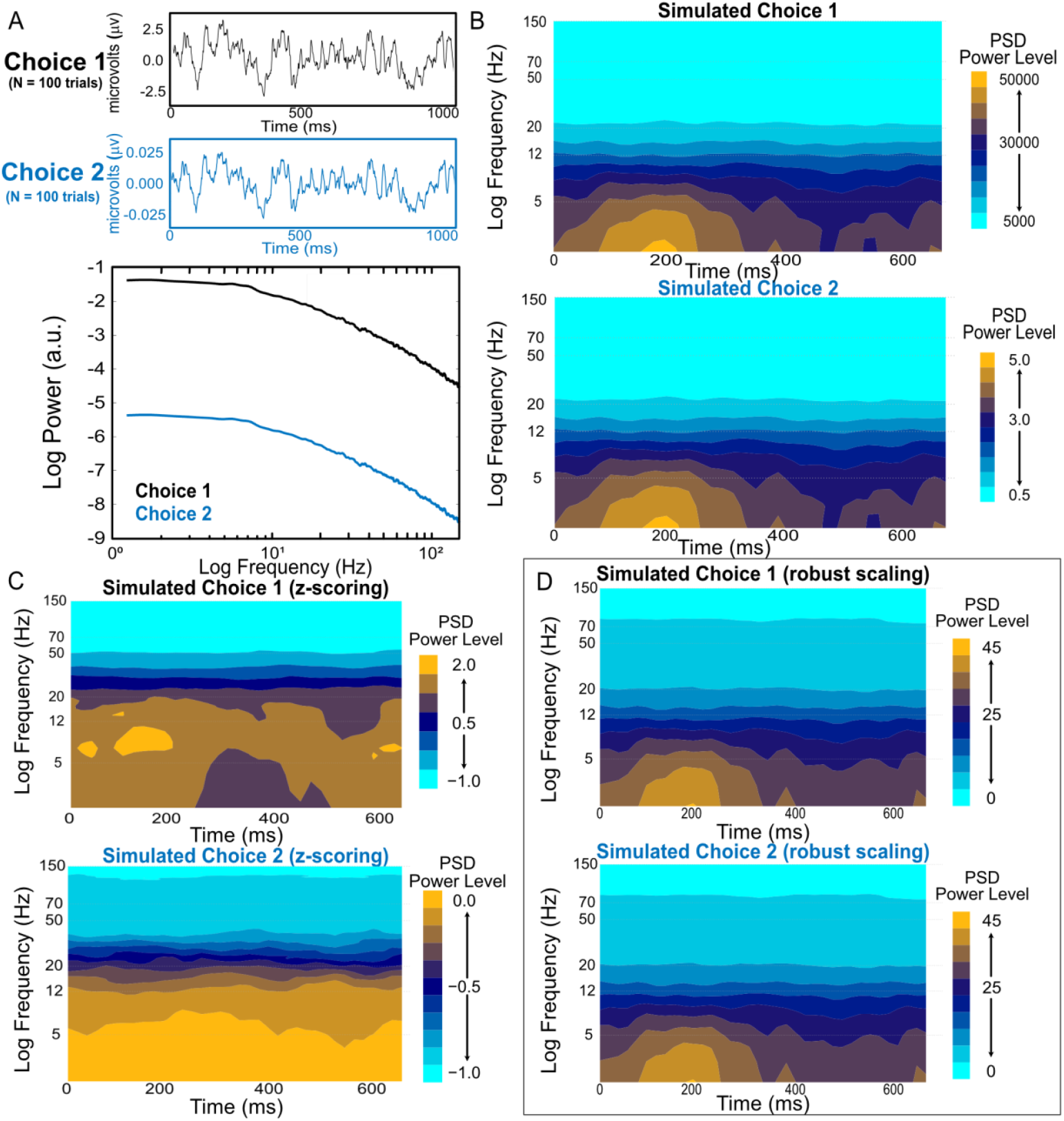
Verification of normalization methods for LFP power by simulation. **A)** A sample LFP trial trace for each choice is shown respectively, and we simulated 100 trials for each choice using the NeuroDSP toolbox. The two choices’ averaged power spectral density (PSD) traces are shown in the log-log regime. The only difference between the two choices was the mean power of PSD across all frequencies. **B)** Prior to normalization, the dynamic of LFP power as a function of frequency and time is shown. The patterns of the two choices looked identical but choice 1’s PSD power levels were 10,000 times greater in amplitude than choice 2’s. **C)** Normalization by balanced z-scoring failed to bring the two choices’ distributions of LFP power to the same scale, and PSD power level was distinctly different across two choices. **D)** Robust scaling succeeded in normalizing LFP power. Note that the two choices shared the same scale of LFP power levels, and the LFP-based CPs computed based on these two choices were 0.5 over frequency and time.

## Acknowledgements

This work was supported by a grant from the Canadian Institutes of Health Research (PJT178071) to CCP. YSH was supported by the China Scholarships Council. We thank Baptiste Caziot and Pooya Laamerad for helpful comments on the manuscript and Julie Coursol for outstanding technical assistance.

## Author Contributions

Y.S.H: formal analysis, visualization, writing – original draft L.D.L: data acquisition, methodology C.C.P: conceptualization, funding acquisition, writing – review & editing

## Data and Code Availability

All original code has been deposited at https://github.com/yueyuesapphirehou/LFP_choice_probability.git and is publicly available. Data reported in this paper will be shared by the corresponding author upon request.

